# Dose-response modelling of total haemoglobin mass to hypoxic dose in elite speed skaters

**DOI:** 10.1101/2020.06.18.159269

**Authors:** Mikhail Vinogradov, Irina Zelenkova

**Affiliations:** Russian Olympic Committee Innovation Center, Moscow, Russia; University of Zaragoza, Spain; Sechenov First Moscow State Medical University (Sechenov University)

**Author notes:** ^#^Corresponding Author: Irina Zelenkova, 125057 Leningradsky av. 75A-62,Moscow, Russia.

**Keywords:** Olympic athletes, simulation, speed skating, total haemoglobin mass response, hypoxic quantification

## Abstract

The aim of the present study is the modelling of the total haemoglobin mass responses in altitude environment with the dose-response model in elite endurance athletes and comparison different existing approaches in the quantification of hypoxic dose.

Data from seven healthy elite endurance athletes specialised in middle distance speed skating participated in the study: six males (24±1.8 years, 182 ±0.3 cm, 84 ±1.5 kg, BMI 23.2±0.6 kg/m^2^, 59.3±1.5 ml/kg/min) and one female (21 years, 164 cm, 56 kg, BMI 17.1 kg/m^2^, 59.9 ml/kg/min). Data were collected during a 3-month training period which included two training camps (14 +14 days) at sea level and two training camps (21+21 days) at altitude of 1224 m and 1850 m above sea level. Total haemoglobin mass (tHb-mass) were measured before the start of the season (baseline) and before and after each training camp (seven measurements) using an optimized CO-rebreathing method, training loads and oxygen saturation at altitude were measured and hypoxic dose were calculated.

Mean total haemoglobin mass for the male group at the base line were 1067±83 g, before the training camp 1 were 1095±82 g, after TC1 1113±105 g, before the training camp 2 (TC2) 1107±88 g, after TC2 1138±104 g. For the female athlete at the base line were 570 g, after TC1 564 g, after TC2 582 g.

The increase of tHb-mass after TC2 were 3,25% and were significant (p<0,005). Mean hypoxic dose for the male group TC1 were %·h (98%) 1078±157, %·h (95%) 79±57, and km.h 473±1 and at TC2 were %·h(98%) 1586±585, %·h (95%) 422±182, and km.h 893±18 and were different from TC1 (p<0,05) for %·h (95%) and km.h methods. For the female athlete hypoxic dose at TC1 were %·h (98%) 970, %·h (95%) 32, and km.h 470 and at TC2 were %·h(98%) 1587, %·h (95%) 289, and km.h 900.

The relationship between hypoxic dose and haematological response was analysed with a non-linear model. The magnitude of the increase of the total haemoglobin mass were investigated using simulation procedures based upon individual responses to the hypoxic dose. We introduced a measurement error to the list square method as a way of avoiding overfitting problem. Dose-response mathematical model between hypoxic dose and total haemoglobin mass was developed. Modelled total haemoglobin mass was within measurement error range. This model is suitable for the computer simulations. The individual response to hypoxic dose due to model data was different. Maximal values in total haemoglobin mass that can be achieved by male athletes according to the model was 1321.9 ± 32 g. The model predicted that (*τ*) erythrocyte life span is 73.8 ± 9.0 days. Moreover, highest value of individual tHb-mass increase after returning to the sea level according to the model was16.3 ±0.7 days.

The model developed in the current study describes the time course of total haemoglobin mass during altitude exposure and post-altitude decline in elite speed skaters.

## Introduction

Altitude training is becoming more and more popular among elite athletes. With innovations in technology and hypoxic methods, a new era of altitude training is currently developing. The 1968 Summer Olympic Games in Mexico City at an altitude of 2250 m attracted the interest of exercise physiologist and elicited new research into altitude physiology. Multiple studies have related the application of altitude training to the improvement of both altitude and sea-level athletic performance (Bonetti, Hopkins, & Kilding, 2006; Faiss et al., 2013; Hahn and Gore, 2001; Katayama, Matsuo, Ishida, Mori, & Miyamura, 2003; Levine and Stray-Gundersen, 1997; Gregoire P. Millet, Roels, Schmitt, Woorons, & Richalet, 2010; J. P. Morton and Cable, 2005; Iñigo Mujika, Sharma, & Stellingwerff, 2019; Schmitt et al., 2006; Wilber, Stray-Gundersen, & Levine, 2007).

Different hypoxic protocols based upon long- and short-term passive exposure, or in combination with different exercise intensities have been described (Girard, Brocherie, & Millet, 2017; Wilber, et al., 2007). Several physiological benefits of altitude exposure are also described in the literature in association with sea-level performance, such as: haematological adaptations, enhanced glycolysis and buffering capacity, increased oxidative capacity, fibre-selective vasodilation and others (Faiss, et al., 2013; Girard, et al., 2017; C. J. Gore et al., 2001; Christopher J Gore et al., 2006; Gregoire P. Millet, et al., 2010; J. P. Morton and Cable, 2005).

In terms of haematological adaptations previous literature has clearly shown that the responses to the hypoxic exposure are very individual and depend upon several factors, including type of altitude (hypobaric/normobaric), altitude itself, the duration of exposure, and others (Bergeron et al., 2012). The measurement of total haemoglobin mass (tHb-mass) has become a ‘gold standard’ of erythropoietic response to altitude exposure (Schmidt and Prommer, 2005). Based upon this measurement a lot of data is now available regarding the individual response to altitude training. Hypoxic exposure may be beneficial for some athletes, but it may also have no effect or be detrimental for others athletes, resulting in lack of mean tHb-mass changes in a group of athletes. Different studies provided different recommendations for hypoxic exposure for athletes to have an increment of tHb-mass. Summarizing these studies, 12 – 16 h.d^−1^ for at least 3 weeks at altitudes 2000 – 2500 m is recommended to increase tHb-mass (Rusko, Tikkanen, & Peltonen, 2004; Wilber, 2007; Wilber, et al., 2007). Based on the available literature on this topic a mean tHb-mass response of ~1% per 100 h of exposure was calculated by Garvican et al. (2012). However this study represented just the very beginning of the discussion on the quantification of hypoxic load and calculation of hypoxic dose. Some formulas were derived and discussions sparked around the approach proposed by Garvican-Lewis and co-workers, where the hypoxic dose was termed “kilometer hours” and defined as km·h = (m/1,000) × h, where m indicates elevation of exposure in meters and h indicates total duration of exposure in hours. (Laura Anne Garvican-Lewis, Sharpe, & Gore, 2016). Other authors, however, questioned the relevance of the quantification of only the “external” stress and proposed a new metric based on “hypoxic stimulus”, termed “saturation hours” and defined as %·h = (98/s − 1) × *h* × 100, where *s* is the saturation value (in %) and h the time (in hours) sustained at any second level (Grégoire P Millet et al., 2016). The above discussion suggests that hypoxic exposure – “external” – stress leads to the “internal” stress, and this impulse can be quantified and described by way of a dose-response model.

Dose-response modelling in sports was introduced by E. Banister and colleagues (E. Banister, Calvert, Savage, & Bach, 1975; Calvert, Banister, Savage, & Bach, 1976). The idea was to use transfer functions to connect input (training load) and output (performance). Training load was mainly described by training impulses (G. P. Millet et al., 2002; G. P. Millet, Groslambert, Barbier, Rouillon, & Candau, 2005; R. H. Morton, Fitz-Clarke, & Banister, 1990), and competition results were used as performance indicators for various sports activities including distance running (Wood, Hayter, Rowbottom, & Stewart, 2005), swimming (I. Mujika et al., 1996; L. Thomas, Mujika, & Busso, 2008), hammer throwing (Thierry Busso, Candau, & Lacour, 1994), and weight lifting (T. Busso et al., 1990; T. Busso et al., 1992)). Attempts have been made to relate several physiological variables (e.g. serum testosterone concentration, cortisol, sex hormone binding globulin, testosterone:cortisol ratio, haemoglobin concentration, red blood cell number, iron status) with fitness or fatigue components of dose-response models (E. W. Banister and Hamilton, 1985; T. Busso, et al., 1990; T. Busso, et al., 1992, Candau, Busso, Lacour, 1992).

More, recently researchers have started to use physiological variables as a model output (Williams et al., 2018). Autonomic nervous system responses were used akin to a performance for dose-response modelling. In principle, it is possible to use such approach to connect any input and output variables. A least square algorithm helps to fit models’ parameters and specify transfer function (Taha and Thomas, 2003).

The main purpose of this study was to build a dose-response model connecting hypoxic dose and total haemoglobin mass. The second goal was to compare different methods of the hypoxic dose quantification.

## Materials and methods

### Participants

Seven healthy elite endurance athletes specialised in middle distance speed skating participated in the study: six males (24±1.8 years, 182 ±0.3 cm, 84 ±1.5 kg, BMI 23.2±0.6 kg/m^2^, 59.3±1.5 ml/kg/min) and one female (21 years, 164 cm, 56 kg, BMI 17.1 kg/m^2^, 59.9 ml/kg/min). All athletes participated in the study during all year (including the competitive season), underwent regular anti-doping controls by international laboratories and were injury and illness free during the study. At the beginning of the study seven males and eight females were recruited to the study but eight participants were excluded from the data analysis (one male and seven females) due to the lack of sufficient data (tHb-mass tests) to perform dose-response modelling. Despite the sample decreased due to organisational issues athlete who left in the study are a small cohort of international level athletes including two world record holders, Olympic bronze medallist and world cup prizewinners. All participants provided written informed consent after being informed of the procedures and risks of involvement in the study. This research was conducted according to the Declaration of Helsinki (2013) and approved by the by the human ethics committee of the Saint Petersburg state university.

### Study design

The study was carried out during the basic period of preparation which included two training camps (mesocycles) at altitude, separated with one mesocycle at sea level. Camp 1: 16 days (Courmayer 1224 m above sea level); 9 days at home; Camp 2: 14 days (Herenveen); 6 days at home; Camp 3: 14 days Minsk; Camp 4: 21 days (Font Romeu 1850 m above sea level). All the athletes underwent a training program according to the preparation phase and trained at least 24 h.wk−1. All athletes trained at the altitude at the first training camp (1200 m) and at the fourth training camp (from 1850 - 2000 m above sea level). Regular control of serum iron was performed and it did not drop below 20 μmol/L throughout the study. In addition, serum ferritin was never indicative of iron deficiency (sFer>20 μmol/L). During altitude exposure daily iron supplementation (100 mg elemental iron with vitamin C 500 mg) was performed.

### Input quantification (hypoxic exposure)

Hypoxic «dose» was calculated using two methods. The first method was based upon the «saturation hours» index and second on «kilometer hours». We measured blood oxygen saturation (SpO_2_) every 2 hours during the day, before bedtime and after waking up, using fingertip pulse oximeters (Choicemmed MD 300 C318, Beijing Choice Electronic Tech Co., Ltd, Beijing, People’s Republic of China). Additionally, during training, blood oxygenation was measured continuously, except during some strengths training exercises where it was technically impossible. Collected data were analysed and «saturation hours» index were calculated using the approach previously described as «saturation hours» (Grégoire P Millet, et al., 2016):

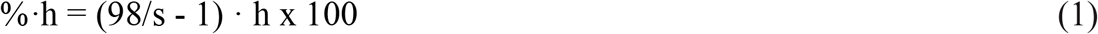

%·h, «saturation hours»; % – arterial oxygen saturation, h-hours of hypoxic exposure.

Hypoxic exposure was also calculated as follows (Laura Anne Garvican-Lewis, et al., 2016):

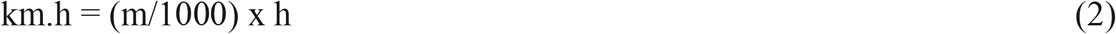

km.h, «kilometer hours» ; km – altitude above sea level, h-hours of hypoxic exposure.

Output quantification (Total haemoglobin mass)

We measured tHb-mass before and after each training camp (seven points) using an optimized carbon monoxide (CO)-rebreathing method (Schmidt and Prommer, 2005). The CO dose was 0.8 mL/kg body mass for female athletes and 1.0 mL/kg for male athletes. The rebreathing procedure was performed for 2 minutes through a glass spirometer (Blood tec GbR, Bayreuth, Germany). Right before the test fingertip capillary blood was taken (200 μl) were collected and at minute 6 and 8 after the start of CO rebreathing (sodium-heparinized 200 μl safe Clinitubes Capillary tubes, Siemens Healthcare Diagnostics Inc., Deerfield, IL, USA) and additionally at minute 6 and 8 after the start of CO rebreathing for determination of %HbCO using ABL80 FLEX CO-OX analyzer (Radiometer, Copenhagen, Denmark). All measurements of tHb-mass were performed by the same researcher using the same equipment with a typical error of 2.0%. Measurements were performed during the rest day at in the laboratory conditions in the morning to avoid any influence of training load on Hb and Htc parameters. Additionally, before the test of tHb-mass concentration Hb (g/dL) and Hct (%) were determined using fingertip capillary blood samples with a blood analyser (Hemo Control; (EKF Diagnostics, Frankfurt, Germany).

### Mathematical model

The model proposed by (Busso, Carasso, & Lacour, 1991) was applied to describe relationships between total haemoglobin mass and hypoxic doses over time. Briefly, this model defines total haemoglobin mass as a sum of the basal level and accumulated hypoxic dose which decay exponentially:

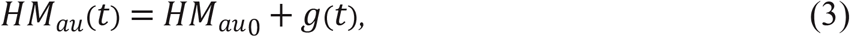

*HM_au_* (*t*) – total haemoglobin mass at day *t* (arbitrary units); *HM*_*au*0_ – basal level of total haemoglobin mass (arbitrary units); *g* (*t*) – positive influence of hypoxic exposure on the haemoglobin mass at day *t*:

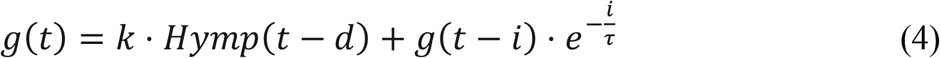

*Hymp* (*t*) is a hypoxic dose at day *t*; *d* – delay of haemoglobin mass response to the hypoxic exposure (days); *k* > 0 is a parameter reflected «strength» of hypoxic influence on the haemoglobin mass; *τ* is a time constant (days), *τ* > 0. The important modification is the existence of the delay parameter (*d*). The hypoxic dose makes a change of the total haemoglobin mass with some time lag.

Usually, Banister’s type dose-response models have two components in transfer function: positive and negative. It is quite natural because of training load influence on the performance in two ways: detrimental and beneficial at the same time. But in the case of the modelling of total haemoglobin mass to hypoxic dose, there is no negative component. The hypoxic exposure influences the total haemoglobin mass only in a positive way.

In the main formula the total haemoglobin mass is expressed in arbitrary units. To calculate actual total haemoglobin mass expressed in grams we used the approach introduced in (Morton, Fitz-Clarke, & Banister, 1990):

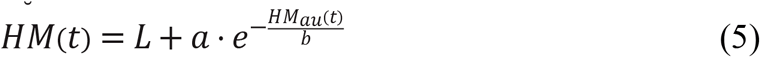

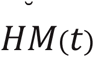 is a total haemoglobin mass at day *t* (grams); *L* is an ultimate limit of the haemoglobin mass for the particular athlete (grams); *a* is a negative amplitude parameter:

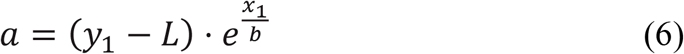

*b* is a positive parameter:

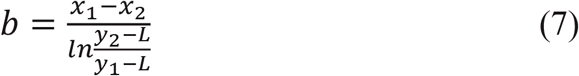

*x*_1_, *x*_2_ are arbitrary points for the two levels of haemoglobin mass (*y*_1_ and *y*_2_)

This new parameter (*L*) takes into account the phenomena of saturation when the athlete is coming to the individual physiological limits of his total haemoglobin mass value.

### Fitting the model

Seven parameters (*t*, *d*, *k*, *L*, *HM*_0_, *a*, *b*) were obtained from the fitting of the modelled haemoglobin mass to its actual values. We used modified least square method, which minimizes the residual sum of squares (RSS) between modelled haemoglobin mass values and

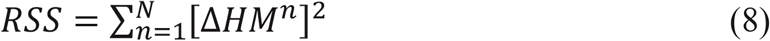

*n* takes the *N* values corresponding to the dates of measurements of the actual haemoglobin mass values (*HM_act_*), and *δ* is the measurement error.

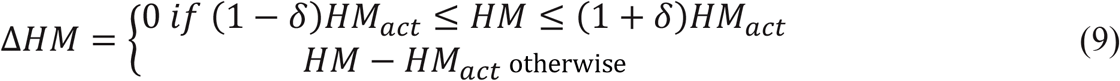

The idea is to prevent the overfitting of the model by allowing for modelled values to stay inside the measurement error interval of the actual values.

### Hypoxic simulation

Simulations of hypoxic exposure were performed for Subject 1. For this participant all simulations began with 9 days of normoxic conditions and then several scenarios were applied. First set of simulations investigated the influence of various altitude exposures with the same accumulated total hypoxic dose. And the second set of simulations were aimed to compare hypoxic expose effect with or without a break (the accumulated total hypoxic dose was the same again).

### Statistics

Statistical analysis was performed using Prism 8 for Mac (GraphPad, USA). Data are presented as mean values ± standard deviation for presented athletes. The typical error (TE) of the tHb-mass measurement were calculated in 10 duplicated measurements with a 24-h gap between the tests dividing the standard deviation of the difference score by root2. To test for significant differences in tHb-mass measured and tHb-mass modelled, Student’s paired t-test was used. P<0.05 was considered to represent statistical significance.

### Results

The real values of tHb-mass for each athlete over the time are presented in Table 1. Mean total haemoglobin mass for the male group at the base line were 1067±83 g, before the training camp 1 (TC1) were 1095±82 g, after TC1 1113±105 g, before the training camp 2 (TC2) 1107±88 g, after TC2 1138±104g. The increase of tHb-mass after TC2 were 3,25% and were significant (p<0,005). For the female athlete at the base line were 570 g, after TC1 564 g, after TC2 582 g.

**Table 1.** Values of the absolute values of tHb-mass (g).

|  | Base line | Before TC1 | After TC1 | Before TC2 | After TC2 |
| --- | --- | --- | --- | --- | --- |
| **Subject 1** | 1129 | 1204 | 1241 | 1210 | 1256 |
| **Subject 2** | 1159 | 1140 | 1176 | 1153 | 1192 |
| **Subject 3** | 1087 | 1103 | 1138 | 1140 |  |
| **Subject 4** | 985 | 1024 | 1019 | 1021 | 1069 |
| **Subject 5** | 976 | 1005 | 993 | 1009 | 1033 |
| **Subject 6** | 897 |  |  | 914 | 935 |
| **Subject 7** | 570 |  | 564 | - | 582 |

Hypoxic dose across all groups using all methods are presented in Table 2.

**Table 2.** Values of different methods of the hypoxic dose quantification.

|  | h(98%) | h(95%) | km.h | h(98%) | h(95%) | km.h |
| --- | --- | --- | --- | --- | --- | --- |
| **Subject 1** | 1321 | 180 | 471 | 1908 | 514 | 856 |
| **Subject 2** | 1107 | 23 | 473 | 1735 | 326 | 901 |
| **Subject 3** | 1003 | 89 | 473 | 518 | 518 | 900 |
| **Subject 4** | 1005 | 65 | 473 | 2172 | 661 | 901 |
| **Subject 5** | 866 | 29 | 473 | 1364 | 140 | 901 |
| **Subject 6** | 1167 | 92 | 473 | 1821 | 376 | 900 |
| **Subject 7** | 1321 | 180 | 471 | 1908 | 514 | 856 |

Mean hypoxic dose for the male group TC1 were %·h (98%) 1078±157, %·h (95%) 79±57, km.h 473±1 and at TC2 were %·h(98%) 1586±585, %·h (95%) 422±182, km.h 893±18 and were different from TC1 (p<0,05) for %·h (95%) and km.h methods. For the female athlete hypoxic dose at TC1 were %·h (98%) 970, %·h (95%) 32, and km.h 470 and at TC2 were %·h(98%) 1587, %·h (95%) 289, and km.h 900. In the beginning we did not find differences between two methods of quantification of hypoxic dose (km-h versus saturation-hours) for the doseresponse relationships if we compare the original equations: km.h = (m/1000) × h %·and h = (98/s − 1) × h × 100. Both methods lead to the same patterns and model parameters except *k*. *k* reflects the strength of HM response for the particular hypoxic dose. For example, for the representative Subject 1 this parameter for the hypoxic impulses quantified as km-hours is estimated at the value of 0.185502341. And in the case of using saturation-hours the parameter is equal to 0.075911079.

After we performed modelling using «saturation hours» with SpO2 98% we faced the problem that despite some athletes from our group receiving hypoxic «dose», we did not see the response in the total haemoglobin mass values. The model with the value 98% wasn’t able to perform any prediction due to absence of response in the face of hypoxic dose input. The question raised related to proposed dose calculation using SpO2 98% and our next step was to compare «saturation hours» with SpO2 98% and 95%.

According to our data there is no difference between two ways of quantification of «hypoxic dose» if we use oxygen saturation 98%. But according to the literature PaOlevels of ∼70 mmHg – is a physiological threshold for the acceleration of the erythropoiesis (Weil, et al., 1968). Also, in the publication of Chapman et al., 2013 they demonstrated a significant difference in EPO with the mean arterial oxygen saturation 95% at 48 collegiate distance runners. At the mean arterial saturation 97% significant difference was not found (R. F. Chapman et al., 2014).

If we recalculate partial pressure in oxygen saturation using the equation developed by Severinghaus (Severinghaus, 1979):

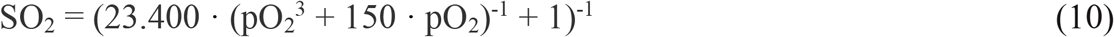

We will receive SaO2 93.8%. According to the medical literature SpO2 95-100% is clarified as a normal value and 91-94% is clarified as a mild hypoxemia (Bledsoe, Porter, Cherry, & Snyder, 2009).

We hypothesized if we use in our model a lower saturation in the «saturation hours» formula we will see difference in these two approaches and we agree with Gregoire M and colleagues that the approach to use the saturation seems to help individualise more the hypoxic dose, especially in athletes with long-term altitude experience.

Thereafter in our model we changed SpO2 98% for 95% in the equation: h = (95/s − 1) × h × 100. In this case, we first of all found a difference between the two approaches of quantification of hypoxic dose (km-h vs saturation-hours). Further results are related to the equation h = (95/s − 1) × h × 100.

Modified formula is:

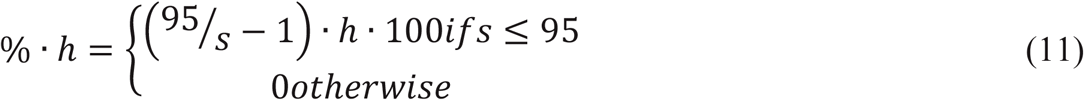

The mathematical model of dose-response relationship between hypoxic impulses (in case of using km-h and sat-hours-95% methods) and total haemoglobin mass fitted real data and the difference between km.h and the model «saturation hours (95%)» were demonstrated. RSS equalled 0 for all seven participants in the case of saturation hours (95%) quantification. RSS equalled 0 only for 5 male athletes. Tables 3–10 present the parameters estimated by the model.

**Table 3.** Main parameter estimates for the proposed dose-response model (male participants, n=6, *HYMPS* are measured in «saturations hours»

| Parameter | $k$ (a.u.) | $\tau$ (days) | $d$ (days) | $HM_0$ (g) |
| --- | --- | --- | --- | --- |
| Mean ± SD | 0,799823 ± 0,7475923 | 72.8 ± 25.8 | 16.0 ± 2.0 | 1151.6 ± 105.1 |
| CV(%) | 93 | 35 | 13 | 10 |

$k$ , multiplying factor for the positive component of hypoxic dose; $\tau$ , time constant of decay for the positive component of hypoxic dose; $d$ , time lag in response of haemoglobin mass to the hypoxic dose; $HM_0$ , basal level of the haemoglobin mass.

**Table 4.** Parameter estimates for the transformation of the criterion haemoglobin mass into the real haemoglobin mass (male participants, n=6, *HYMPS* are measured in «saturations hours»).

| Parameter | $a$ (a.u.) | $b$ (a.u.) | $L$ (g) |
| --- | --- | --- | --- |
| Mean ± SD | $-351.22 \pm 216.10$ | $684.56 \pm 561.57$ | $1321.92 \pm 78.32$ |
| CV(%) | -62 | 82 | 6 |

$a$ , magnitude parameter in equation converting haemoglobin mass in arbitrary units to haemoglobin mass in grams.; $b$ , the denominator of the exponent in above equation; $L$ , an ultimate limit of the haemoglobin mass for the particular athlete.

**Table 5.**
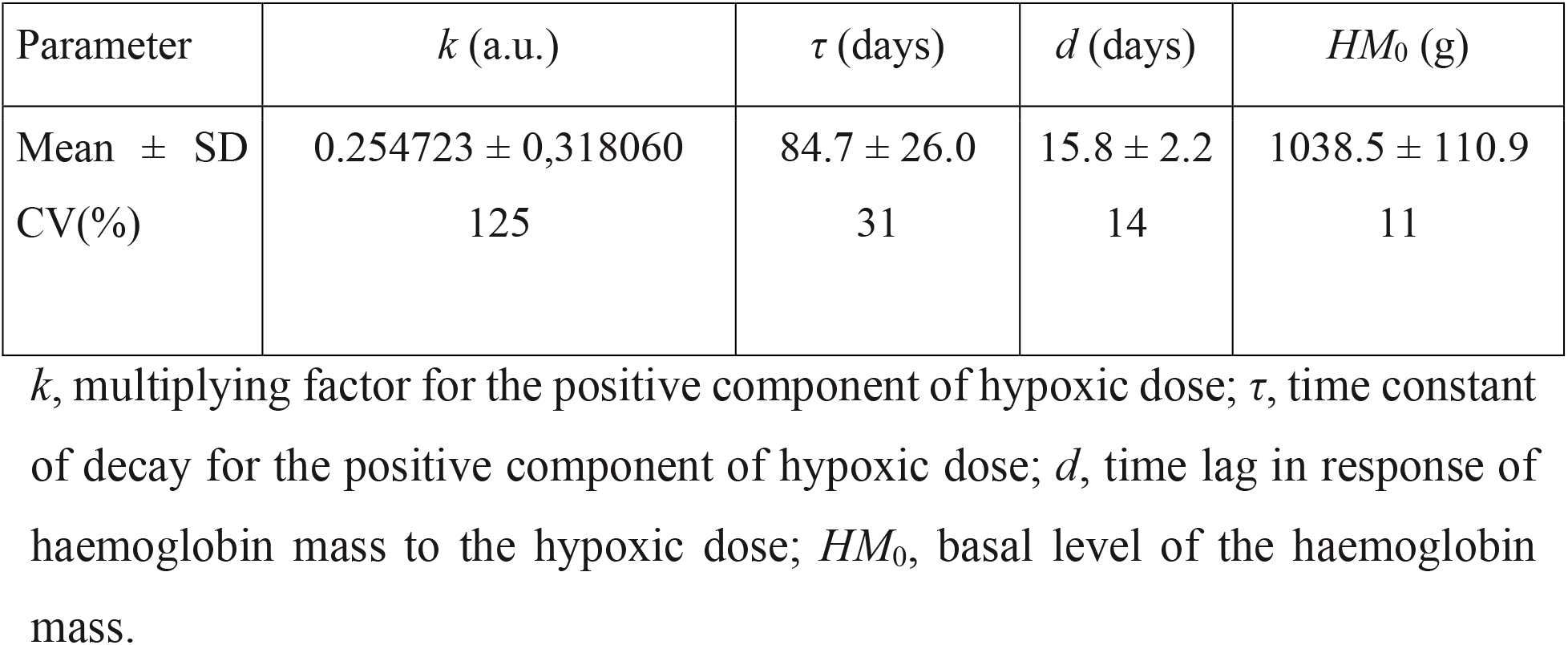
Main parameter estimates for the proposed dose-response model (male participants, n=5, *HYMPS* are measured in «km-hours»)

| Parameter | $k$ (a.u.) | $\tau$ (days) | $d$ (days) | $HM_0$ (g) |
| --- | --- | --- | --- | --- |
| Mean ± SD | $0.254723 \pm 0,318060$ | $84.7 \pm 26.0$ | $15.8 \pm 2.2$ | $1038.5 \pm 110.9$ |
| CV(%) | 125 | 31 | 14 | 11 |

$k$ , multiplying factor for the positive component of hypoxic dose; $\tau$ , time constant of decay for the positive component of hypoxic dose; $d$ , time lag in response of haemoglobin mass to the hypoxic dose; $HM_0$ , basal level of the haemoglobin mass.

**Table 6.**
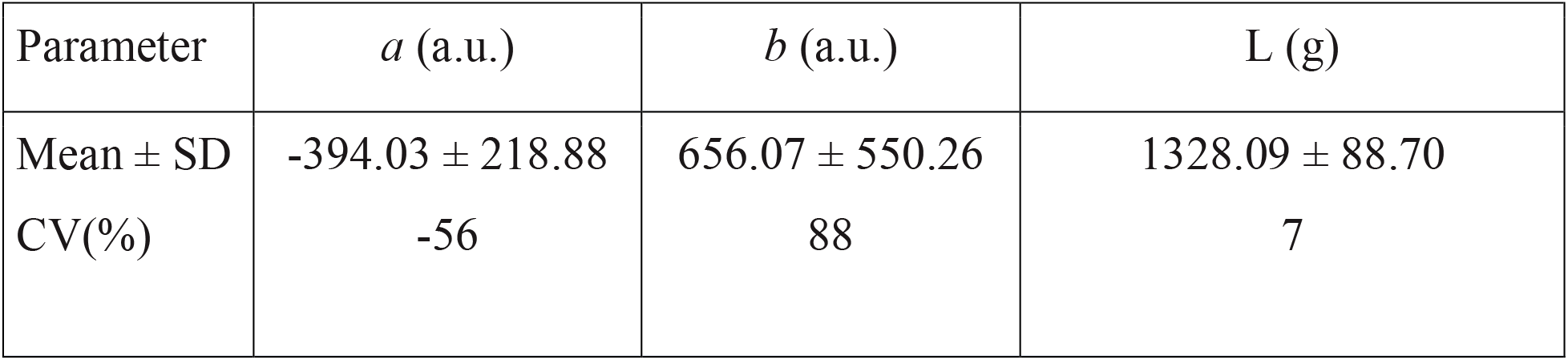
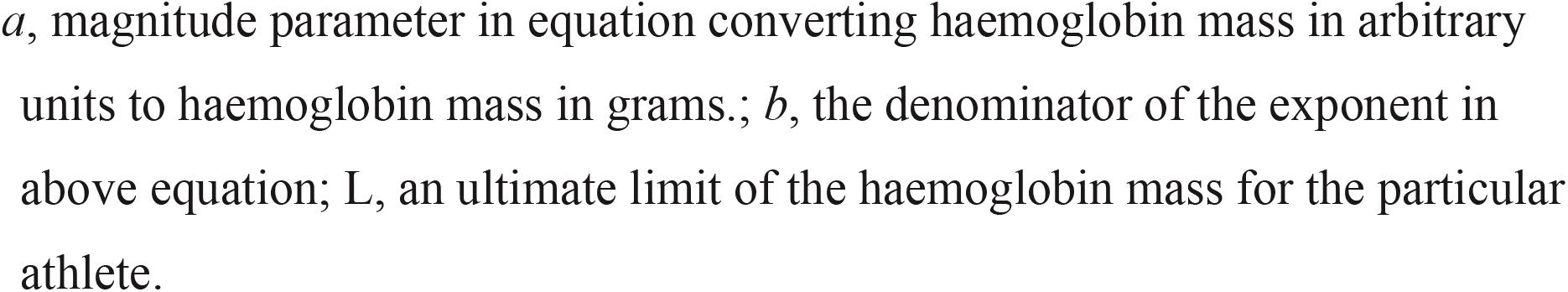
Parameter estimates for the transformation of the criterion haemoglobin mass into the real haemoglobin mass (male participants, n = 5, *HYMPS* are measured in «km-hours»)

| Parameter | $a$ (a.u.) | $b$ (a.u.) | $L$ (g) |
| --- | --- | --- | --- |
| Mean ± SD | $-394.03 \pm 218.88$ | $656.07 \pm 550.26$ | $1328.09 \pm 88.70$ |
| CV(%) | -56 | 88 | 7 |

$a$ , magnitude parameter in equation converting haemoglobin mass in arbitrary units to haemoglobin mass in grams.; $b$ , the denominator of the exponent in above equation; $L$ , an ultimate limit of the haemoglobin mass for the particular athlete.

**Table 7.**
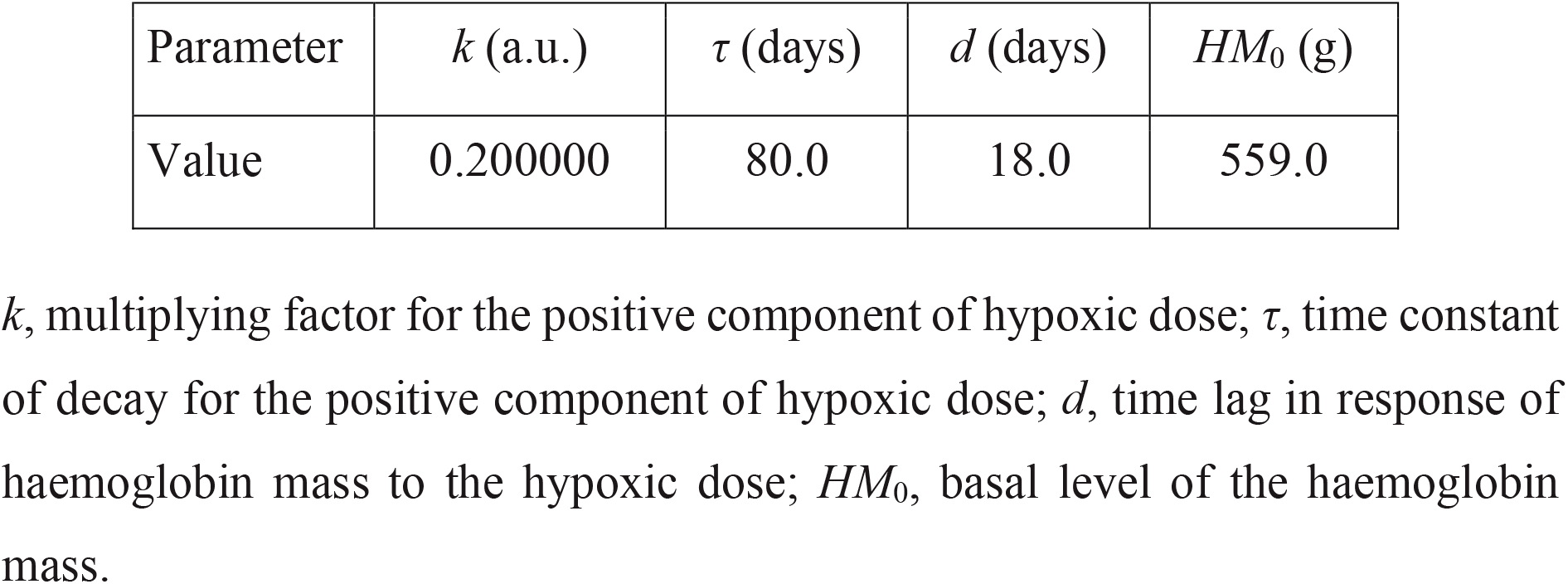
Main parameter estimates for the proposed dose-response model (female participant, n=1, *HYMPS* are measured in «saturations hours»):

**Table 8.** Parameter estimates for the transformation of the criterion haemoglobin mass into the real haemoglobin mass (female participant, n=1, *HYMPS* are measured in «saturations hours»):

| Parameter | $a$ (a.u.) | $b$ (a.u.) | $L$ (g) |
| --- | --- | --- | --- |
| Value | -489.69 | 409.12 | 942.50 |

$a$ , magnitude parameter in equation converting haemoglobin mass in arbitrary units to haemoglobin mass in grams.; $b$ , the denominator of the exponent in above equation; $L$ , an ultimate limit of the haemoglobin mass for the particular athlete.

**Table 9.** Main parameter estimates for the proposed dose-response model (female participant, n=1, *HYMPS* are measured in «km-hours»):

| Parameter | $k$ (a.u.) | $\tau$ (days) | $d$ (days) | $HM_0$ (g) |
| --- | --- | --- | --- | --- |
| Value | 0.03 | 25.0 | 18.0 | 565.0 |

$k$ , multiplying factor for the positive component of hypoxic dose; $\tau$ , time constant of decay for the positive component of hypoxic dose; $d$ , time lag in response of haemoglobin mass to the hypoxic dose; $HM_0$ , basal level of the haemoglobin mass.

**Table 10.** Parameter estimates for the transformation of the criterion haemoglobin mass into the real haemoglobin mass (female participant, n=1, *HYMPS* are measured in «km-hours»):

| Parameter | $a$ (a.u.) | $b$ (a.u.) | $L$ (g) |
| --- | --- | --- | --- |
| Value | -481.18 | 412.08 | 942.50 |

$a$ , magnitude parameter in equation converting haemoglobin mass in arbitrary units to haemoglobin mass in grams.; $b$ , the denominator of the exponent in above equation; $L$ , an ultimate limit of the haemoglobin mass for the particular athlete.

The dynamics of haemoglobin mass and hypoxic dose (%h and km hours) are presented in Figure 1 (a,b)

**Figure 1.**
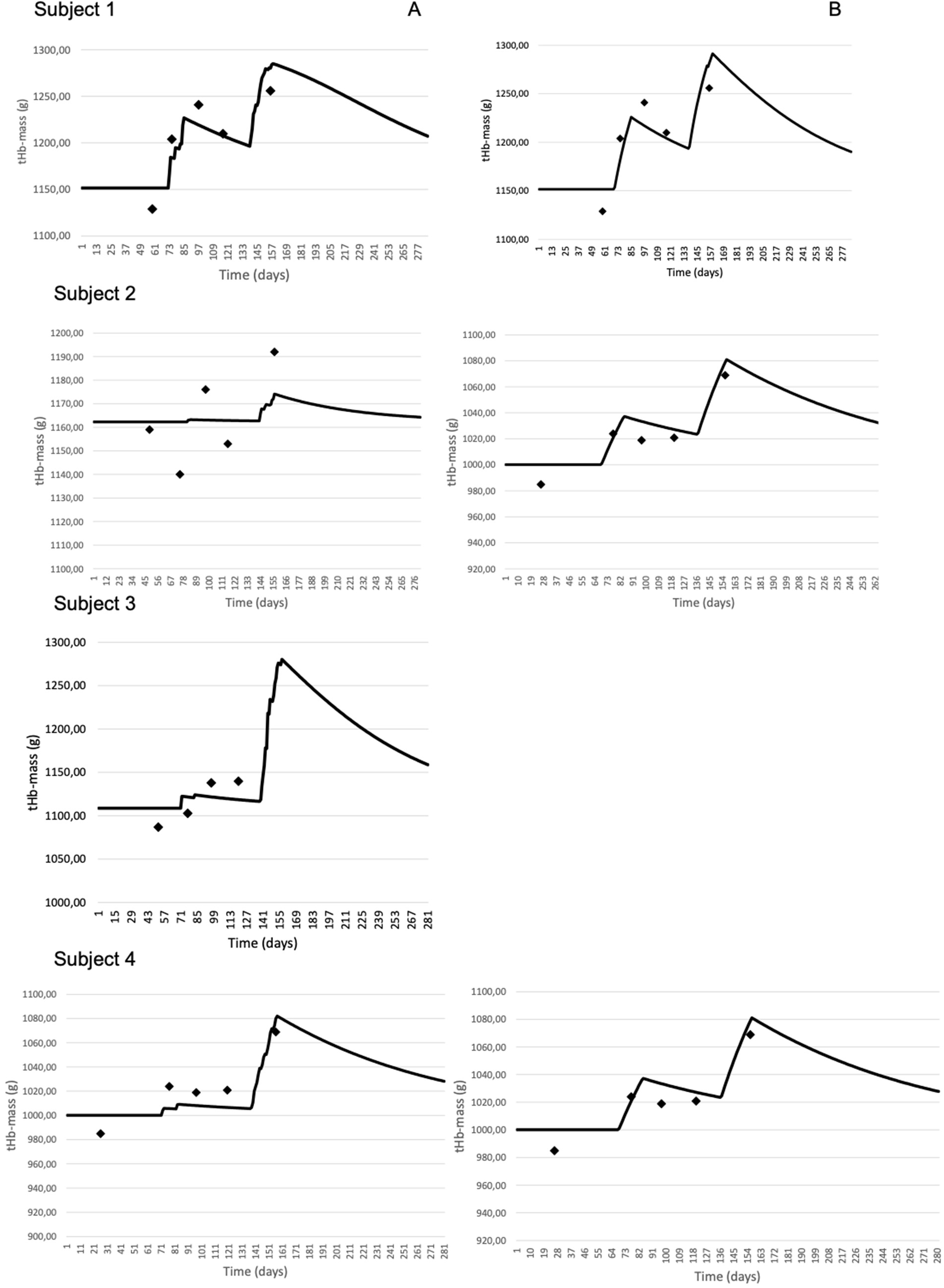
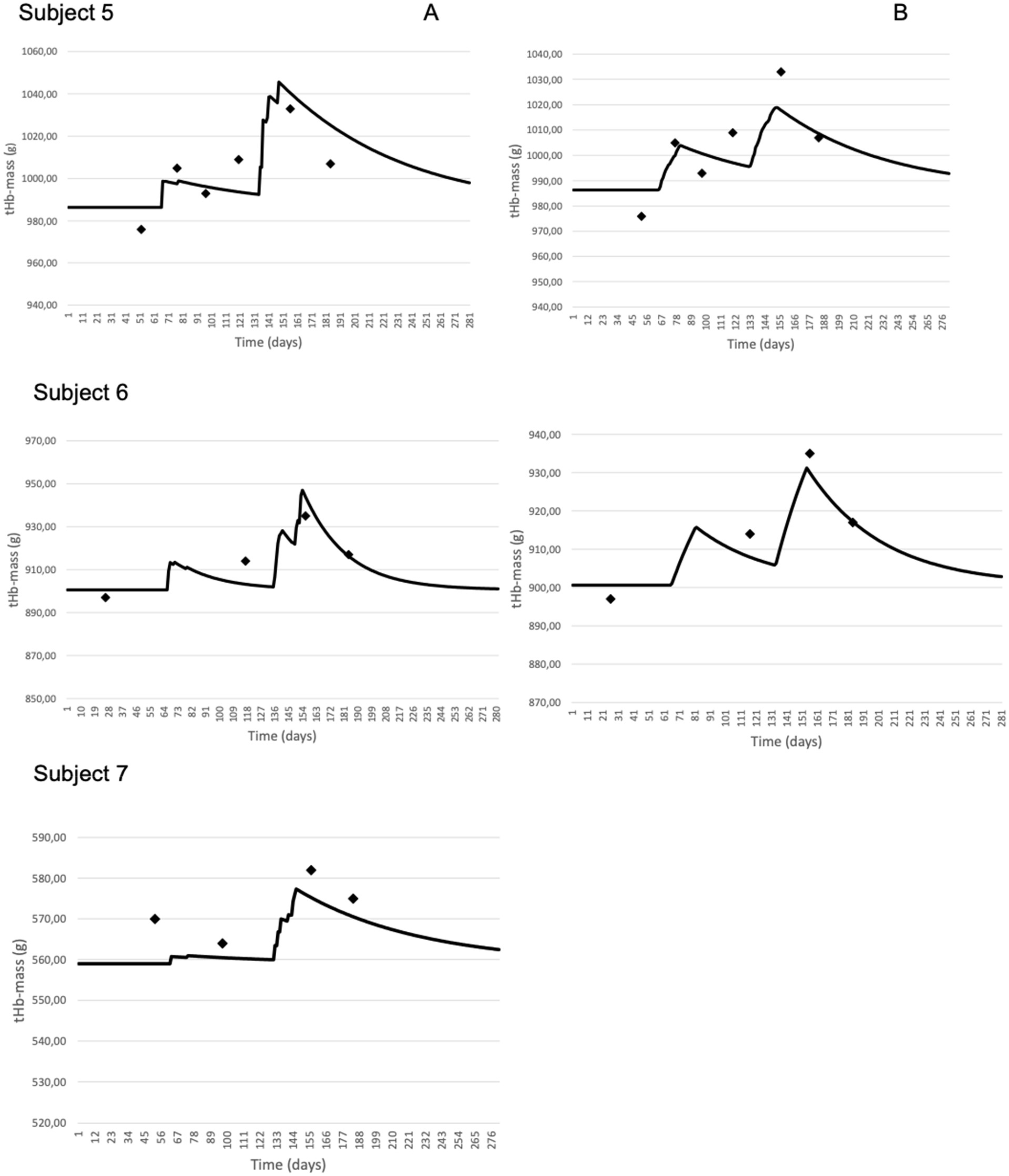
Application of the model. Measured (points) and modelled (line) haemoglobin mass for %h (95) (A) and km hours (B).

**Figure 2.**
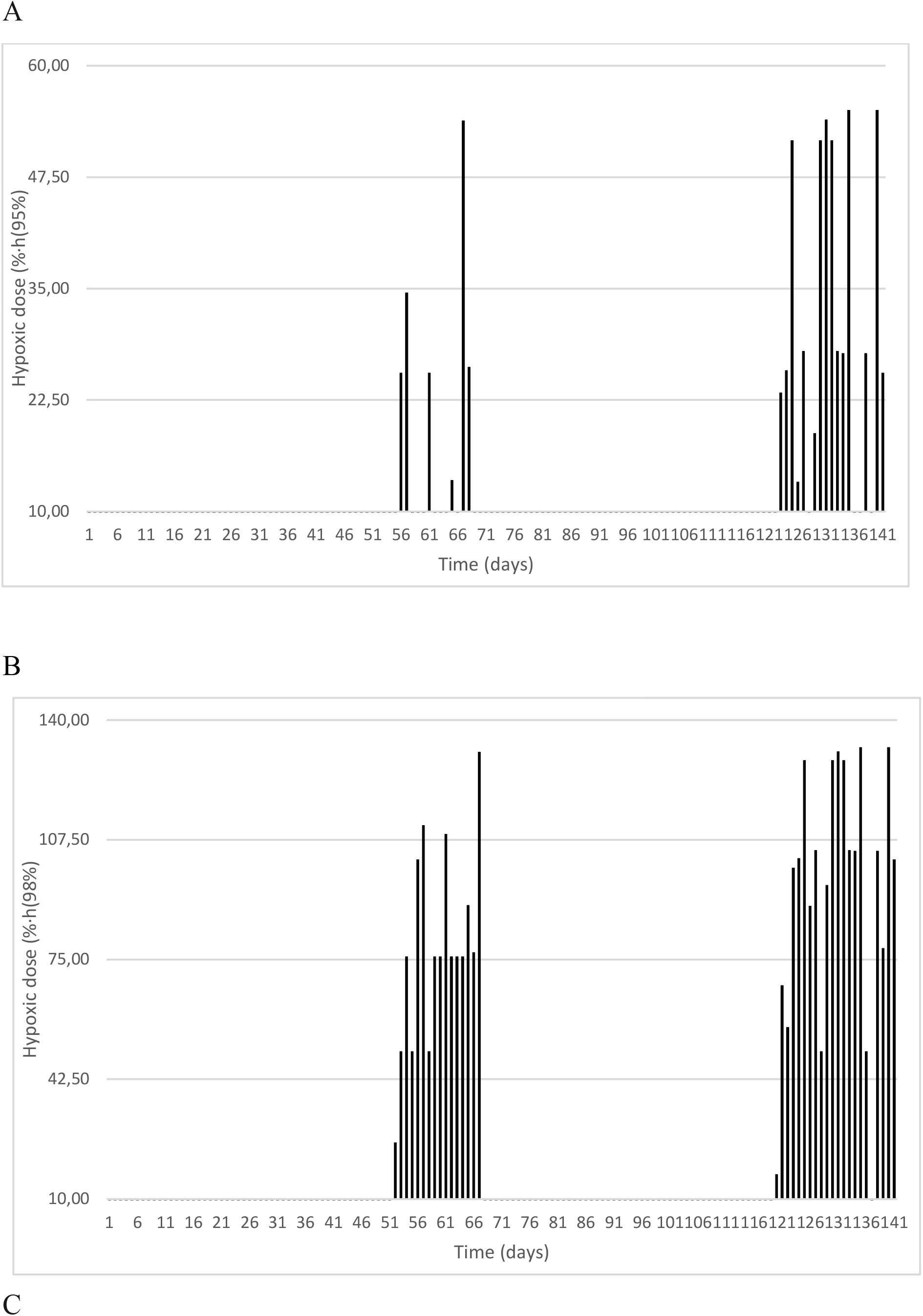
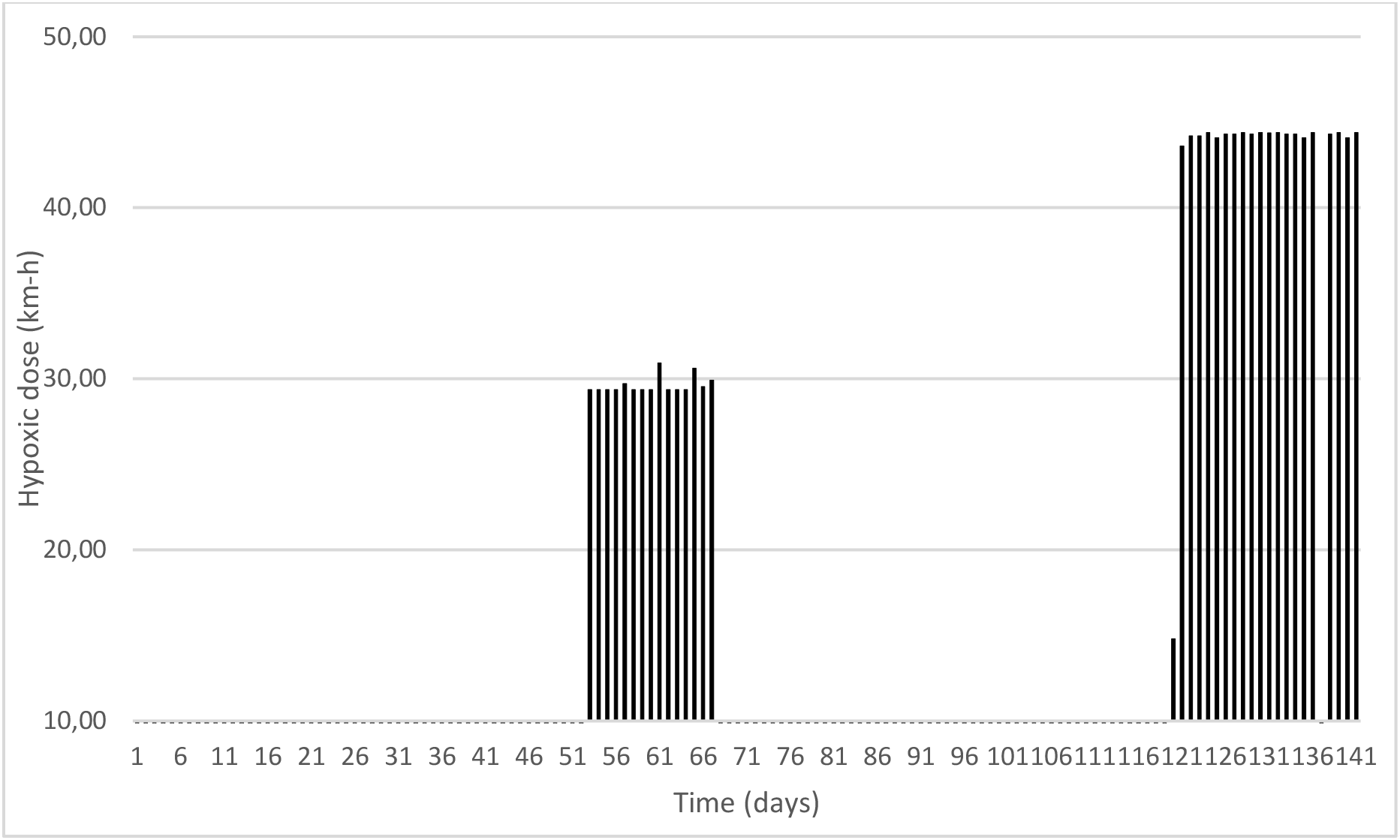
Hypoxic dose measured in % h(95%) (A), % h(98%) (B) and km hours (km-hours) (C)

After performing Student t-test we didn’t find any difference between the modelled absolute tHb-mass values and the real data.

### Comparison of two methods of quantification of hypoxic impulses

According to our data, the model based upon km.h approach does not fit the real data. In contrast the model based upon «saturation hours» (95%) fitted perfectly the real data.

**Table 3.** Comparison of modelled absolute tHb-mass (g) and real data.

|  | Mod-<br>elled<br>(BL) | Differ-<br>ence,<br>% | Mod-<br>elled<br>BTC1 | Differ-<br>ence,% | Mod-<br>elled<br>ATC1 | Differ-<br>ence,<br>% | Mod-<br>elled<br>ATC2 | Differ-<br>ence,<br>% | After<br>TC2 | Mo<br>del<br>led | Differ-<br>ence,<br>% |
| --- | --- | --- | --- | --- | --- | --- | --- | --- | --- | --- | --- |
| **Subject 1** | 1152 | 2.0 | 1184 | -1,7 | 1219 | -1.8 | 1208 | -0.2 | 1256 | 12<br>80 | 1.9 |
| **Subject 2** | 1154 | -0,5 | 1163 | 2.0 | 1172 | -0.3 | 1169 | 1.4 | 1192 | 11<br>94 | 0.2 |
| **Subject 3** | 1109 | 2.0 | 1122 | 1,7 | 1122 | -1.4 | 1118 | -1.9 |  |  |  |
| **Subject 4** | 1000 | 1.5 | 1021 | -0,3 | 1033 | 1.4 | 1027 | 0.6 | 1069 | 10<br>79 | 0.9 |
| **Subject 5** | 986 | 1.1 | 1000 | -0,5 | 1001 | 0.8 | 997 | -1.2 | 1033 | 10<br>18 | -1.5 |
| **Subject 6** | 901 | 0.4 |  |  |  |  | 908 | -0.7 | 935 | 93<br>0 | -0.5 |
| **Subject 7** | 559 | -1.9 |  |  | 560 | -0.6 |  |  | 582 | 57<br>5 | -1.2 |

### Simulation of various combination of the hypoxic exposure

Additionally, in our study using «dose-response» modelling we tested several scenarios with the same hypoxic dose (1512 km-hours) but different elevation and duration (Fig 3 A,B):

**Fig 3A.**
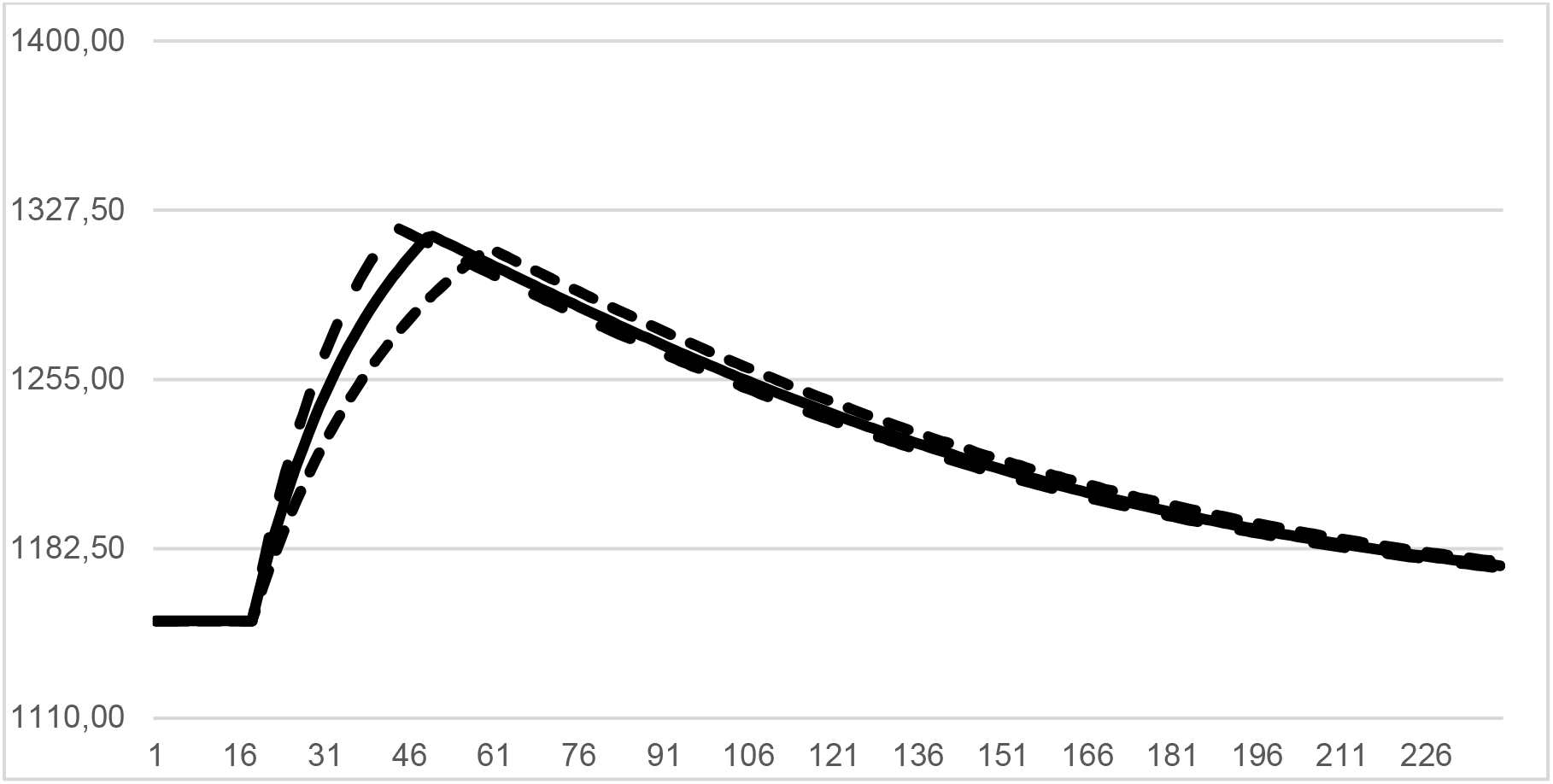
Simulations obtained from data and model parameters of a representative athlete. Simulation: I – 42 days at the altitude 1500 meters (short dash line); II – 31.5 days at the altitude 2000 meters (solid line); III – 25.2 days at the altitude 2500 meters (long dash line).

**Fig 3B.**
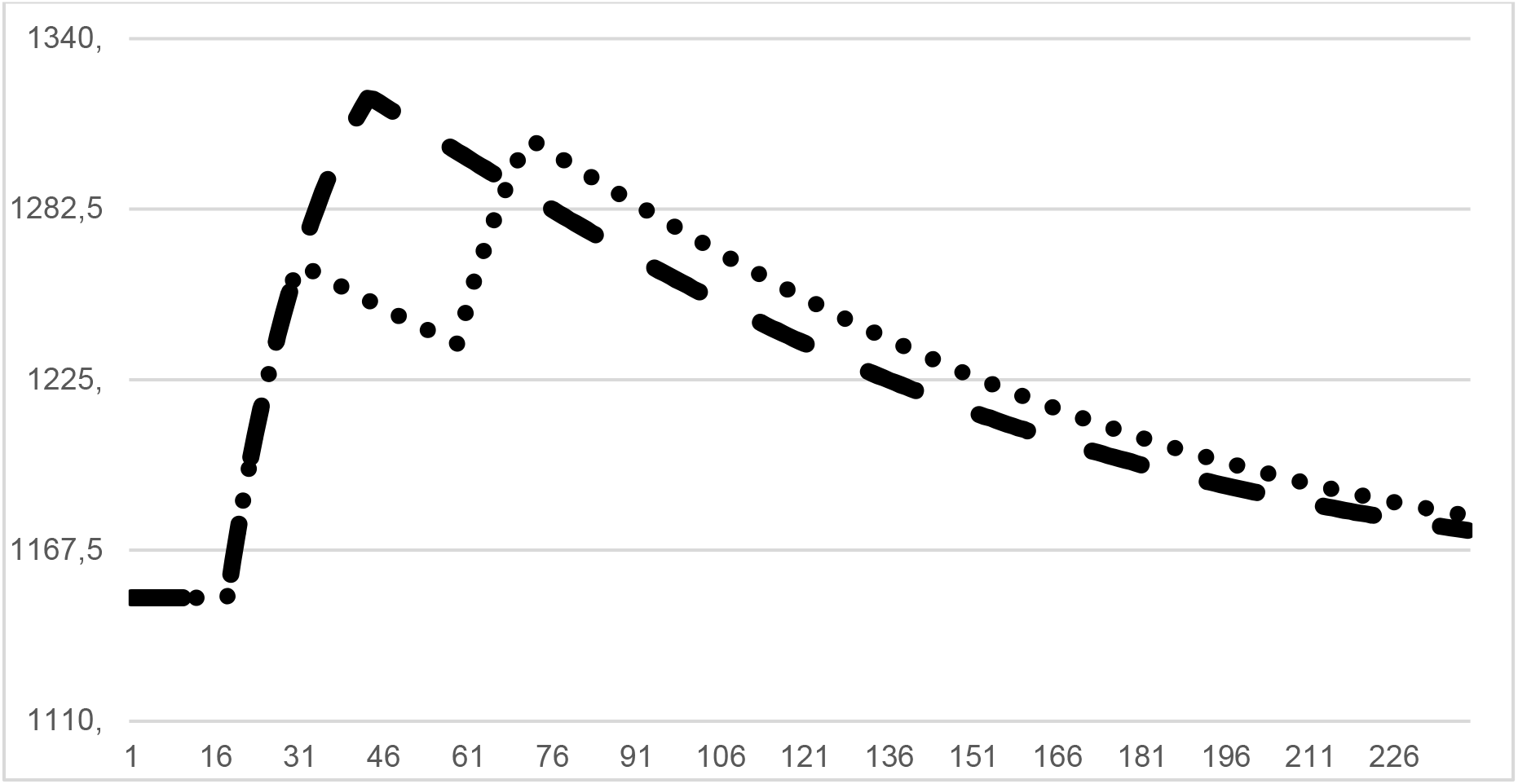
Simulations obtained from data and model parameters of a representative athlete. Simulation: I – 12.6 days at 2000 m, 28.4 days at sea level and 12.6 days at 2000 meters (dotted line); II – 25.2 days at the altitude 2500 meters (solid line). Peak tHb mass is different in each scenario: I – 1306.09 g, II – 1319.92 g

The tendency is clear, but, the difference between various scenarios is within the measurement error (2%). So, the total accumulated hypoxic dose (without breaks) is preferable according to the tHb mass gain.

According to the modelling results we could see that same hypoxic dose received in different time frame leads to different patterns of tHb mass increase that can be helpful for athletes for the year planification and pre-competitive tapering.

The individual response to hypoxic dose due to model data was different (*k*) and was 0.7 ± 0.3. Maximal values in total haemoglobin mass that can be achieved by male athletes are 1321.9 ± 32 g. The model predicted that (*tau*) erythrocyte life span is 73.8 ± 9.0 days. Moreover, optimal performance after training at the moderate altitude were 16.3 ± 0.7 days.

The developed model in the current study describes time course of total haemoglobin mass during the altitude exposure and post-altitude effect in elite speed skaters.

The peak level of tHb-mass is achieved at the 18th day after the cessation of hypoxic exposure. This time pattern is influenced by the lag parameter of the model (*d*).

## Discussion

Despite the debates that are still ongoing behind the effects of altitude training athletes and coaches, especially in endurance sports use hypoxic exposure in their preparation (Iñigo Mujika, et al., 2019).

Since the concept of the utilisation of altitude training in preparation for the performance at altitude as a pre-acclimatisation, other concepts for the increase of sea-level performance have been developed. The approach of the Living High – train at Low altitude and High altitude seems to be one of the most effective due to different physiological effects associated with performance enhancement: aerobic and anaerobic (Christopher John Gore, Clark, & Saunders, 2007).

Variation in oxygen level in the blood is important mechanism in the adaptation of the body to the altitude. Despite that altitude training has a very complex impact on the body and leads to numerous adaptations such as haematological, cardiovascular, ventilatory, improvement in muscle buffer capacity, glycogen enzyme activity, muscle mitochondrial volume and others that leads to performance enhancement in elite athletes haematological adaptations seems to be the most important in the concept of altitude training (Otto et al., 2017; Zelenkova, Zotkin, Korneev, Koprov, & Grushin, 2019). Increase of aerobic performance seems to be related with a signalling mechanism associated with the tissue-dependent up-regulation of HIF-1a and HIF-2a, increase of EPO secretion and followed increase in tHb-mass and blood volume and thereby increase in VO2max (Lundby, Calbet, & Robach, 2009; Schmidt and Prommer, 2008) that allows to deliver more oxygen to the working muscles (Levine and Stray-Gundersen, 2005). This signalling pathway we took as a basis of our model. Our study was carried out on elite athletes at the moderate altitude (1850-2400 m) that is regular used by sport practitioners to enhance altitude and sea level performance and a lot of data from the literature are available (R. Chapman and Levine, 2007; Flaherty, O’Connor, & Johnston, 2016). When an athlete faces the hypoxic exposure, the first trigger that starts the adaptation process is decrease of the arterial oxygen saturation (SaO2) that leads to the increase of the EPO production by the kidneys (Jelkmann, 2011). According to the literature the partial pressure of oxygen ∼70 mmHg is enough to accelerate erythropoiesis (Weil, Jamieson, Brown, & Grover, 1968). After arrival at the altitude EPO level increases and according to several studies reaches the peak at 1-3 day (Clark et al., 2009; Jelkmann, 2011).

Erythropoietic response is a very individual parameter due to the lack of EPO measurements in our study we were not able to use time course of EPO in our model (R. F. Chapman, Stray-Gundersen, & Levine, 1998).

After going through the cell proliferation from the early erythroid progenitor cell differentiation after several days the cell becomes a mature blood cell. The life span of red blood cells may vary from 70 to 140 days and the mean lifespan is 115 days that falls in line with our model (Robach et al., 2006).

Across the wide range of studies related to the altitude training, individual response to the hypoxic exposure in athletes can be even opposite: from significant increase in some athletes and even decrease in others (Płoszczyca, Langfort, & Czuba, 2018). In our study, time course of the total haemoglobin mass is in line with the study of Garvican who documented the measurable increase of tHb-mass of 3% at the 11th day after the start of altitude training (Garvican, et al., 2012). According to the model, following 3 weeks of continuous exposure to 1850 meters, the EPO response increases tHb-mass by 3.7 % or 43.4 g (or 0.52 g per kg). Such an increase in tHb-mass, according to the publications, should increase absolute VO2max by 17.2 mL/min in well-trained athletes (Schmidt and Prommer, 2008) free of any influence of the factors listed above that can decrease erythropoiesis. These results are in line with the study with the same altitude exposure that was carried out by Garvican-Lewis et al., were after 21 days of altitude exposure at 1850 meters we observed increase in 3.0% (Laura A Garvican-Lewis, Halliday, Abbiss, Saunders, & Gore, 2015). The increase demonstrated in our study is also close to the magnitude observed in previous studies with the same hypoxic exposure 1% for every 100 h above 2100 meters (C. J. Gore et al., 2013). The main difference in our study is the exposure altitude. The present study was carried out in the natural altitude of 1850 meters and athletes had at least 23 hours of hypoxic exposure every day during three weeks accumulating the following sufficient amount of hours in hypoxia: 161 h after 7 days, 322 h after 14 days, 483 h after 21 day (Christopher J Gore and Hopkins, 2005). According to the linear model with the dose expressed as «kilometer х hours» that predicts the increase of approximately 3.4 % with “hypoxic dose” ranged from ~890 km.h to ~1400 km.h. In our study the increase was 3,25% and according to the model 3.7%.

One of most important topics in the practice is the time-course of haemoglobin mass after return to the sea level and the maintenance of the post altitude training effects. In the model we focused on the increase of the tHb-mass. Despite this there are publications that elucidate the negative impact of return to the sea level on the tHb. According to the study of Garvican et al., where they observed a 1.5% decrease in tHb-mass after 3 days of descent from altitude and which persisted when measured 10 days after descent (Garvican, et al., 2012). This decrease in RBS is selective and mostly premature (young) RBCs are subjected to the neocytolysis and selective hemolysis of circulating red blood cells (Alfrey, Rice, Udden, & Driscoll, 1997). Mechanisms behind this process is most probably associated with the down-regulation of HIF-controlled BINIP3L – regulated autophagy along with a decrease in RBC antioxidant catalase (CAT) in erythrocytes produced after hypoxic exposure and mitochondrial-derived reactive oxygen species (ROS) in reticulocytes (Song, Sundar, Gangaraju, & Prchal, 2017). According to the publications different drops of total haemoglobin mass are described: in the study of Prommer et al., that was performed on a group of elite Kenyan runners after descending from the altitude 2090 meters, haemoglobin mass started to decrease after 14 days at sea level (Prommer et al., 2010). Seems that it depends from the duration of the altitude exposure and altitude. But as we see from several studies total haemoglobin mass after 1 or 2 weeks after the return to the sea level, the mean increase is around 8% (Garvican, et al., 2012; Prommer, et al., 2010). This fact shows that neocytolysis plays the minor role in post altitude red cell mass volume change and can be neglected or the parameters for the model should be more sensible to see the impact of the reticulocyte loss.

According to the study of Garvican et al., on return to sea level at day 10, mean tHb-mass remains stable D31 2.3 (0.6-4.1)% and was still «likely higher» comparing to the baseline (Garvican, et al., 2012). In other studies there was documented significantly higher level of tHb-mass comparing to the base level after 21 day of the return to the sea level (Brocherie et al., 2015) that also was in line with the estimation of the model proposed by Gore et al., where higher levels of tHb-mass are estimated for 21 day (C. J. Gore, et al., 2013). Our model is in line with this estimation where we see very gradual drop of tHb-mass. More research is required to confirm or disprove these estimations.

This impressive difference across the athletes in tHb-mass change during altitude training can be associated with several factors such as: iron/vitamin deficiency, inflammation/injury, erythropoietic response to the altitude (Friedmann et al., 2005). But one of the main parameters that we consider important is individual hypoxic «dose» that every athlete receives during the altitude exposure and quantification of hypoxic impulse remains to be one of the principle information that elucidates the magnitude in individual response. Physiological response to hypoxia varies from subject to subject, and arterial oxygen saturation seems to be more accurate in predicting individual response to the hypoxic-induced haematological response. In our study we compared two approaches: «kilometer hours» and «saturation hours».

In our study amongst the 7 athlete for whom we performed simulations we found that model km.h did not fit two athletes’ data and the model based on «saturation hours» fitted all the data. But from a practical point of view for the planification and routine practice «km hours» seems to be easier to use. Despite during our study we observed individual response in athletes of the saturation related to the training intensity these small oscillations did not influence the full picture of the general altitude exposure.

The dose-response model in our study allows us to make simulation and observe effects of various scenarios of hypoxic dose and time frame of hypoxic exposure. The idea of using dose-response models for simulations is popular in the research papers. The main focus was in the simulation of various taper scenario. The study of triathlete shows the result that progressive taper is beneficial then step (E. W. Banister, Carter, & Zarkadas, 1999). But the 2-phase pattern with a final increase in training leads to an even better outcome (Luc Thomas, Mujika, & Busso, 2009). Optimal duration of taper varies between 2 and 4 weeks (T. Busso and Thomas, 2006). But in case of pre-taper overload training period it should be extended (Sanchez et al., 2013). And simulations allow to conclude that the overload period is necessary to achieve better performance (L. U. C. Thomas and Busso, 2005). Also there is possibility to use simulation to find training load saturation thresholds and predict overtraining phenomena (R. H. Morton, 1997; Turner, Mazzoleni, Little, Sequeira, & Mann, 2017). The ability of the dose-response model to predict the effect of the particular training load or overall training program is attractive to practitioners (T. Busso, Benoit, Bonnefoy, Feasson, & Lacour, 2002; Méline, Mathieu, Borrani, Candau, & Sanchez, 2019). And using simulations from our model are in line with these modelling efforts.

The weak point of the model relates to the athlete individual response. If we have an illness/injury condition, anaemia, small physiological response to the altitude weaker than typical error of the measurement or not enough measurement the model is not able to make any predictions. The limitation of the study is related with the small sample size. Future studies needed with the bigger sample size and different cohorts (regional, national athletes and untrained individuals).

As a conclusion of our discussion we would like to point out that there is crucial importance to unify the «hypoxic dose» metrics and hypoxic dose in kilometer hours, as it seems to be quite accurate and very practical approach for the altitude training guidelines, planification and quantification. The developed model in current study describes time course of total haemoglobin mass during the altitude exposure and post-altitude effect.

In conclusion, we would like to point out that that the effect of altitude training varies from many parameters and it should be very carefully planned and supervised by a specialist in altitude training to avoid mistakes. The proposed model can be very useful tool for the individualisation and planification of the athlete is exposed to altitude and can be for the great interest for coaches and sport scientists.

## Acknowledgements

The authors express deep acknowledgements to the “Russian skating union”. The authors thank all athletes in their commitment to the study and team specialists Poltavets K., Gogotova V., Yakushkin A., Rozina M. The authors acknowledge team staff to their help and support during all the study. Also authors would like to acknowledge Iñigo Mújica, Mike Davidson and Yannis Pitsiladis for their valuable help in the preparation of this manuscript.

